# Temporal attention selectively enhances target features

**DOI:** 10.1101/2021.03.25.437071

**Authors:** Luis D. Ramirez, Joshua J. Foster, Sam Ling

## Abstract

How does directing attention to a moment in time augment vision? Here, we examined the computations by which temporal attention – the allocation of attention to a moment in time – improves perception, under a divisive normalization framework. Under this framework, attention can improve perception of a target signal in three ways: *stimulus enhancement* (increasing gain across all sensory channels), *signal enhancement* (selectively increasing gain in channels that encode the target stimulus), or *external noise exclusion* (reducing the gain in channels that encode irrelevant features). These mechanisms make diverging predictions when a target is embedded in varying levels of noise: stimulus enhancement improves performance only when noise is low, signal enhancement improves performance at all noise intensities, and external noise exclusion improves performance only when noise is high. To date, temporal attention studies have used noise-free displays. Therefore, it is unclear whether temporal attention acts via stimulus enhancement (amplifying both target features and noise) or signal enhancement (selectively amplifying target features) because both mechanisms predict improved performance in the absence of noise. To tease these mechanisms apart, we manipulated temporal attention using an auditory cue while parametrically varying external noise in a fine-orientation discrimination task. Temporal attention improved performance across all noise levels. Formal model comparisons revealed that this cueing effect was best accounted for by a combination of signal enhancement and stimulus enhancement, suggesting that temporal attention improves perceptual performance, in part, by selectively increasing gain for target features.

## Introduction

Our ability to appropriately respond to dynamic environments involves the recruitment of temporal attention, the allocation of attention to a moment in time. A growing body of evidence has demonstrated that temporal attention improves perceptual detection and discriminability (Griffin, Miniussi, and Nobre 2001; Correa, Lupiáñez, and Tudela 2005; Rohenkohl et al. 2012), which is thought to be mediated by improvements in early visual processing (Correa, Lupiáñez, et al. 2006; Correa, Sanabria, et al. 2006; Rolke and Hofmann 2007). However, the computational mechanisms subserving these improvements in target detection and discriminability due to temporal attention remain unclear (Nobre and Rohenkohl 2014).

Discriminating a target stimulus in noise is a classic signal detection problem, where performance is governed by the ratio between the intensity of the signal and the intensity of the noise, both in the environment and in the visual system itself (Pelli and Farell 1999). Within this framework, attention might improve the signal-to-noise ratio in several ways (Lu and Dosher 2008). First, attention could increase the gain of all visual features, amplifying both relevant signal and irrelevant noise (Lu and Dosher 1998; Dosher and Lu 2000b), which we call “stimulus enhancement”. Second, attention could selectively increase the gain of the target signal, leaving any irrelevant noise untouched, which we call “signal enhancement” (see Footnote). Finally, attention could improve visual processing by suppressing irrelevant noise, thereby improving the signal-to-noise ratio, “noise exclusion” (Dosher and Lu 2000b; Dosher et al. 2004).

Stimulus enhancement, signal enhancement, and noise exclusion each have distinct signatures depending on the amount of noise present in a display (Figure 1). Notably, stimulus enhancement and signal enhancement both predict an improvement in perceptual sensitivity in the absence of noise, as has been reported in past studies of temporal attention (Nobre, Correa, and Coull 2007; Nobre and van Ede 2018; Shalev, Nobre, and van Ede 2019). However, because temporal attention studies have used noise-free displays, it is unclear whether temporal attention improve perception via stimulus enhancement or signal enhancement. In this study, we parametrically varied external noise to directly test which mechanism supports temporal attention.

**Figure 1.**
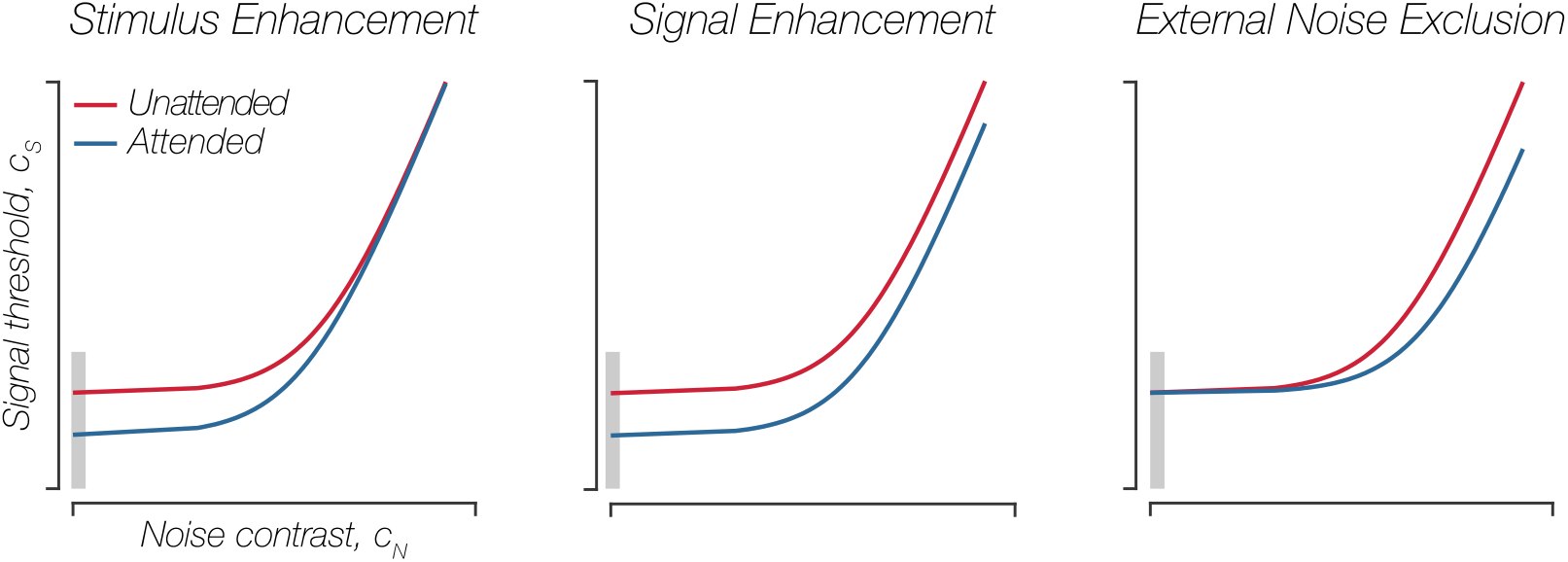
Perceptual Thresholds Under Predicted Attention Mechanisms. Red curves represent perceptual thresholds across increasing levels of noise in the absence of attention. Blue curves represent perceptual thresholds across increasing levels of noise under attention. Note how each attentional mechanism distinctly improves signal contrast thresholds across increasing levels of noise. Stimulus enhancement evokes a threshold reduction primarily at low noise levels, signal enhancement evokes threshold reduction across all noise levels, whereas noise exclusion evokes threshold reduction primarily at high noise levels. Note also that stimulus enhancement and signal enhancement cannot be distinguished in the absence of noise (gray rectangle).

Several studies have manipulated external noise to test how spatial attention modulates perception (Lu and Dosher 1998; Dosher et al. 2004; Lu and Dosher 2008; Ling, Liu, and Carrasco 2009; Pratte et al. 2013). These studies typically used a variant of signal detection models, such as the Perceptual Template Model (PTM), to model signal contrast thresholds as a function of external noise. However, while the PTM can dovetail nicely with behavioral data, it assumes that the effect of external noise is additive with the signal, which recent work has shown is not the case (Baker and Vilidaite 2014; Baldwin, Baker, and Hess 2016; Hansen and Hess 2012). Instead, the effect of noise is better accounted for by gain control mechanisms, where the signal and noise inhibit each other. This mutual suppression is thought to occur due to divisive normalization (Morrone, Burr, and Maffei 1982; Freeman et al. 2002; Brouwer and Heeger 2011; Carandini and Heeger 2012; Ling and Blake 2012). Therefore, we adopted a model in which the interaction between the signal and noise are governed by normalization. Under normalization models, the neural response to an item is determined by the balance between excitatory and inhibitory neural activity. Specifically, the neural response to a stimulus is regulated by its own response, as well as adjacent neural responses (Carandini and Heeger 2012). This framework has long been deployed to account for interactions within visual cortex (Heeger 1992) and has more recently been proposed to play a role in the modulatory effects of attention (Ling and Blake 2012; Bloem and Ling 2019; Ruff and Cohen 2017; Reynolds and Heeger 2009). Within this framework, attention can improve our ability to detect signals in noise by tipping the balance between neural excitation and inhibition. Our variant of the normalization model of attention integrates the predicted mechanisms of attention from perceptual template models to generate distinct, testable hypotheses for how temporal attention enhances perceptual sensitivity: stimulus enhancement, signal enhancement, and noise exclusion (Figure 1).

Under *stimulus enhancement*, attention boosts the neural representation of the stimulus in its entirety — both relevant target signal and irrelevant distractor noise. Thus, stimulus enhancement improves target discrimination primarily when external noise is low because this mechanism also amplifies noise. Under *signal enhancement*, attention solely boosts the target signal, thereby improving target discrimination even when the target is embedded in noise. A final possibility is that temporal attention elicits *external noise exclusion* — reducing the neural representation of noise, primarily when noise is high. However, on its own, noise exclusion cannot explain the finding that temporal attention improves performance in the absence of noise (Nobre, Correa, and Coull 2007; Anna C. Nobre and van Ede 2018; Shalev, Nobre, and van Ede 2019). Nevertheless, we consider external noise exclusion in our model comparisons because temporal attention might evoke external noise exclusion in combination with signal enhancement.

In this study, we combine the predicted mechanisms of attention from perceptual template models with the visual cortical interactions described under normalization models to test whether temporal cues improve visual sensitivity through stimulus enhancement, signal enhancement, noise exclusion, or a combination of mechanisms. Participants performed a fine-orientation discrimination task on a target grating that appeared randomly in time and was masked by white noise whose contrast was parametrically manipulated. In half the trials of this task, participants had no knowledge of the target grating’s onset; this served as our uncued (unattended) condition. This was compared to our cued (attended) condition, where participants were provided an auditory cue that immediately preceded the target grating — providing precise temporal information about the target signal’s impending onset and the moment in time a participant should attend. To assess the effect of the alerting cue on perception, we measured signal contrast thresholds under cued and uncued conditions across multiple contrast levels of the noise mask. We found that temporal attention boosts perceptual sensitivity across all noise levels. Moreover, in an additional experiment, we empirically rule out the possibility that these effects are solely driven by changes in temporal uncertainty from cueing. Taken together, our results suggest that temporal attention improves perception, in part, via signal enhancement, selectively enhancing processing of target features.

## Methods

### Participants

Twelve healthy adult volunteers between ages 18-24 (7 females; age = 20.92 ± 1.14, mean ± standard error of the mean, SEM) participated in the experiment. One subject was not included in data analyses because perceptual thresholds did not increase monotonically with noise contrast and were outliers. An outlier was determined if the average signal contrast threshold, collapsed across noise contrast levels for each condition, was greater than 2.5 standard deviations from the mean.

All participants had normal or corrected-to-normal vision. A minimum sample size of ten was chosen comparable to other studies that have utilized a similar making paradigm (Lu and Dosher 1998; Dosher and Lu 2000a; Lu and Dosher 2000; Dosher et al. 2004). We also ran six participants in a control experiment (see below for more information), including three subjects from the main experiment, and three newly recruited subjects (3 females; age = 27.66 ± 2.11, mean ± SEM). For two of the six subjects in this control experiment, we collected data across the original ten external noise levels in the orientation discrimination task portion of this control experiment and thus include their signal contrast thresholds in the model fitting procedure for the main study. All participants involved provided written consent and were reimbursed for their time. The Boston University Institutional Review Board approved the study.

### Apparatus and stimuli

Stimuli were generated using MATLAB (The Math Works Inc 2007) in conjunction with the Psychophysics Toolbox (Brainard 1997), rendered on a Mac Mini running Ubuntu 16.04 LTS. Stimuli were presented on a gamma-corrected CRT monitor (1280- × 1024-pixel resolution; 75 Hz refresh rate), with no additional light sources in the room. Participants were seated comfortably with their heads in a chin rest at a viewing distance of 57 cm from the screen. The background of the display was uniform gray (luminance = 49 cd/m^2^).

### Task procedure

Participants performed a fine-orientation discrimination task in which they reported the tilt of a target grating embedded in dynamic noise (Figure 2a). We parametrically manipulated the contrast of this noise mask from trial to trial (ten noise contrast levels evenly spaced on a log scale between 0% and 34.66% RMS contrast; Figure 2b). Each trial began with the onset of a dynamic

**Figure 2.**
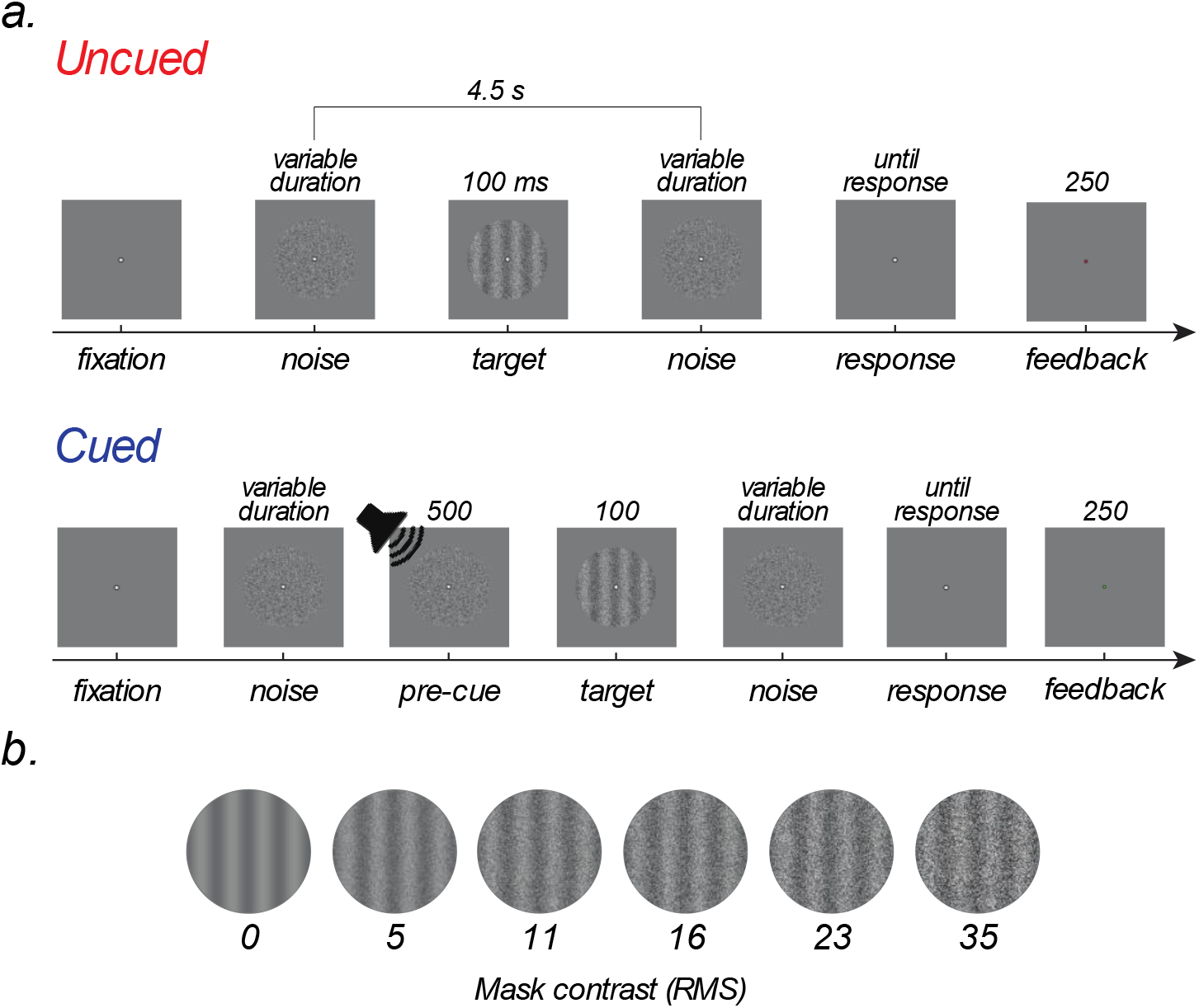
Masking Paradigm and Experimental Design. A. Trial sequence. Each trial began with fixation, followed by the onset of the mask display after 500ms, which remained on for the entirety of the trial (roughly 4500 ms). Within a trial, the target grating randomly appeared for 100ms. In half the trials, this target was immediately preceded by an auditory cue (500ms), providing precise temporal information about the target’s onset. Subjects reported the target grating’s orientation after target offset and were provided feedback after the end of each trial. B. Examples of the experimental stimuli: target gratings masked in various levels of external noise (Gaussian white noise).

Gaussian white noise mask (diameter = 4°, changing at 10 Hz; subtending 0.2° in diameter) at fixation. The noise mask was present for the full duration of the trial (jittered between 4.5–4.7s). Participants were instructed to maintain steady fixation through each trial. The target grating (spatial frequency = 6 cycles/°, fixed spatial phase, diameter = 4°, orientation = ± 2° from vertical) was presented for 100 ms within the noise mask. The target grating could appear at the following timepoints within a trial: 1 s, 1.6 s, 2.8 s, or 4.0 s. Participants had no knowledge of the number of possible timepoints the target could appear at or of how these timepoints were generated. On half of the trials, the participants were presented with an auditory cue that cued when the target would appear (*cued trials*). In these trials, the target grating was preceded by a 100% valid auditory cue that swept from 262 Hz (C_4_) to 880 Hz (A_5_) across a 500-ms period. The target grating immediately followed this cue, drawing participants attention to the exact moment that the target grating would appear. On the other half of trials, no auditory cue was presented (*uncued* trials). Following target offset, participants reported whether the target was tilted clockwise or counterclockwise from vertical. Participants had no limit on response time to emphasize accuracy over response time. Feedback was provided at the end of each trial for 250 ms, followed by a 750-ms inter-trial interval. Feedback consisted of a change in color of the fixation dot from white if the response was correct (green), incorrect (red), or a wrong key press (grey).

Prior to the main blocks of the task, participants completed a training block that contained all conditions randomly interleaved (2 attention conditions × 10 noise mask levels × 2 target orientations). We included this training block to ensure that participants were familiar with the timing of events in each trial. Participants were informed during training that the auditory cue was 100% valid and immediately preceded the target grating.

Participants completed 2–3 sessions of the task in total, where each session consisted of 800 trials (40 trials per condition). Trials in each condition were interleaved with their order randomized in each experimental session. A break was provided every 40 trials. We used an adaptive staircasing procedure, QUEST (Watson and Pelli 1983), to estimate contrast thresholds for the target grating in each condition for a total of 20 independent staircases (2 attentional conditions × 10 noise contrast levels) set to a performance level of 70% accuracy (d’ = 0.74). All staircases operated continuously across sessions, each receiving 40 trials in each session. If a staircase had not converged at the end of a session (operationalized as the standard deviation of the threshold distribution being above 0.1), the subject completed an additional session until all staircases met this criterion. To satisfy this requirement, subjects completed 2-3 sessions, resulting in 80 or 120 trials per condition for each subject.

### Model fitting procedure

To determine which attention mechanism best characterized the observed attention effect, we first fit the reduced normalization model (Equation 1) to each subject’s contrast thresholds from the uncued condition. This model is a modified Naka-Rushton:

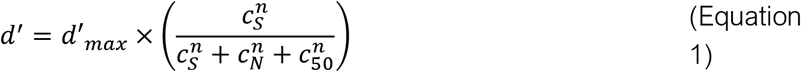

where d′ represents discriminability or perceptual sensitivity, d′_max_, maximum perceptual sensitivity, *c_S_*, contrast of the signal (the target grating), *c_N_*, contrast of the noise mask, *c_50_*, semi-saturation point, and *n*, dynamic range or a non-linear transducer. The parameters that represent attention mechanisms — stimulus enhancement, signal enhancement, and external noise exclusion — are excluded from the reduced model to establish a baseline in the absence of attention. Solving for the observer’s signal contrast threshold in this reduced model generates predicted Threshold vs. Contrast curves (Equation 2)(Blakemore and Campbell 1969).

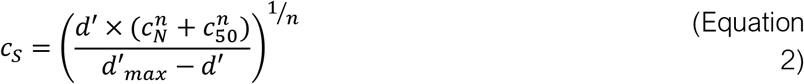

Using nonlinear regression, we fit each subject’s signal contrast thresholds in the uncued condition with the reduced model. Initial parameter values for d′_max_, *c_50_*, and *n* were chosen based on a series of grid searches for the most optimal initial parameter values that generated the lowest sum of squared errors, then estimated using the *fmincon* function in MATLAB. Next, we fit variants of the modified normalization model to the measured signal contrast thresholds. Each variant of the model allowed a different attentional coefficient (or combination of attentional coefficients) to vary while fixing d′_max_, *c_50_*, *n* to the estimated values from the reduced normalization model. The full normalization model, including all attention mechanisms, is expressed as follows (Equation 3):

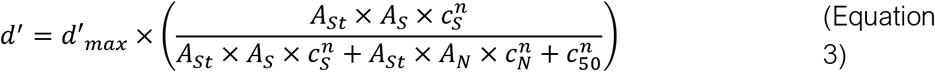

*A_St_* is the stimulus enhancement coefficient, acting on both the signal, *c_S_*, and noise, *c_N_*. *A_S_* is the signal enhancement coefficient, acting solely on the signal. Finally, *A_N_* is the noise exclusion coefficient, acting strictly on the external noise. All attention coefficients were constrained to be between values of 0 and 5, where a value of 0 produces a complete suppression of the response to a stimulus component (signal, *c_S_*, or noise, *c_N_* depending on the attention coefficient), a value of one produces no attentional modulation compared to the reduced model, and values greater than one produce attentional modulation that enhances a stimulus component. Solving for signal contrast thresholds results in the following (Equation 4):

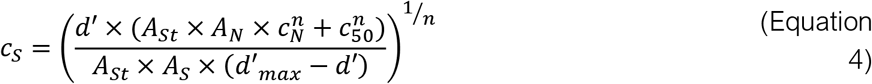

Each attention mechanism and combination of attention mechanisms were accounted for, resulting in a total of six additional variations of the modified normalization model to fit to the cued condition’s data to for each subject, using the *fmincon* function in MATLAB.

To evaluate which mechanisms could most parsimoniously account for our data, we used a corrected version of the Akaike Information Criterion, AICc (Akaike 1974; Cavanaugh 1997). This metric accounts for the number of observations and free parameters in a model to estimate the relative amount of information loss. The lower the AICc value, the better a given model explains the data. If we compute the difference between the minimum AICc value and all AICc values for each subject, we expect that the better a model, the closer to 0 the difference will be on average.

### Control Experiment

We conducted a control experiment to test whether the cuing effect could be explained by decreased temporal uncertainty about when the target grating appeared. Participants (n=6) performed a detection task, in which they reported the presence or absence of a grating. The stimulus parameters and sequence of trial events in the detection task were identical to the orientation discrimination task. The probability of whether the target grating was present or absent on a given trial was drawn from a uniform discrete distribution. As in the orientation discrimination task, the auditory cue was present in half the trials. Participants performed this task across five of the 10 external noise contrasts used in the main experiment (0%, 1.44%, 4.76%, 10.53%, and 34.66% RMS contrast), which spanned the full range of noise contrasts used in that experiment. The contrast of the target grating was set to each participant’s signal contrast thresholds obtained in the orientation discrimination task. Of the six participants, three participants took part in the main experiment. For these participants, signal contrast thresholds were obtained in the main experiment. The remaining participants completed sessions of the fine-orientation discrimination task used in the main experiment until the standard deviation of the signal contrast threshold distribution for each staircase was below 0.1. For two of the six subjects in this control experiment, we collected data across the original ten external noise levels in the orientation discrimination task portion of this control experiment and thus include their signal contrast thresholds in the model fitting procedure for the main study.

All 6 subjects in the control experiment completed at least two sessions of the fine-orientation discrimination task. One subject completed three sessions of the fine orientation discrimination task because their staircases had not yet converged after the second session, meaning the standard deviation of the signal contrast threshold distribution for each staircase was not yet below 0.1 after the second session.

## Results

We found that signal contrast thresholds were lower in the cued condition than in the uncued condition across all levels of noise contrast (Figure 3a). Figure 3b shows the percent increase in signal contrast thresholds between the cued and uncued conditions. To test which mechanism of attention best accounted for the temporal cueing effect across noise levels, we fit a family of normalization models to signal contrast thresholds (see Methods, model fitting procedure). We found that the reduced model (with all attention coefficients set to 1), fit nicely to the uncued data (min & max R^2^: 0.5726, 0.9366; average R^2^: 0.839 ± 0.036; c_50_: 0.063 ± 0.016; **n**: 1.775 ± 0.494; 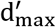:3.405 ± 0.333; mean ± SEM). Unsurprisingly, the baseline (reduced) model, with fixed parameters estimated from the uncued data for each subject, fit the cued data poorly (average R^2^: 0.371 ± 0.101; mean ± SEM), suggesting that an absence of attention mechanisms is insufficient for explaining the cued condition’s data for each subject (Figure 4). The average ΔAICc across all subjects revealed that a combination of stimulus enhancement and signal enhancement is the winning model on average (Figure 5), followed closely by signal enhancement alone (ΔAICc values: A_ST_ & A_S_ = 3.207 ± 0.787; A_S_ = 3.308 ± 1.589; A_ST_ & A_N_ = 6.393 ± 1.770; A_S_ & A_N_ = 6.3939 ± 1.770; A_ST_ = 6.658 ± 2.961; A_N_ = 11.984 ± 2.422; baseline = 12.083 ± 2.853; mean ± SEM). Thus, our results suggest that stimulus enhancement alone does not account for the data best. Instead, our results show that temporal attention recruits signal enhancement in addition to stimulus enhancement.

**Figure 3.**
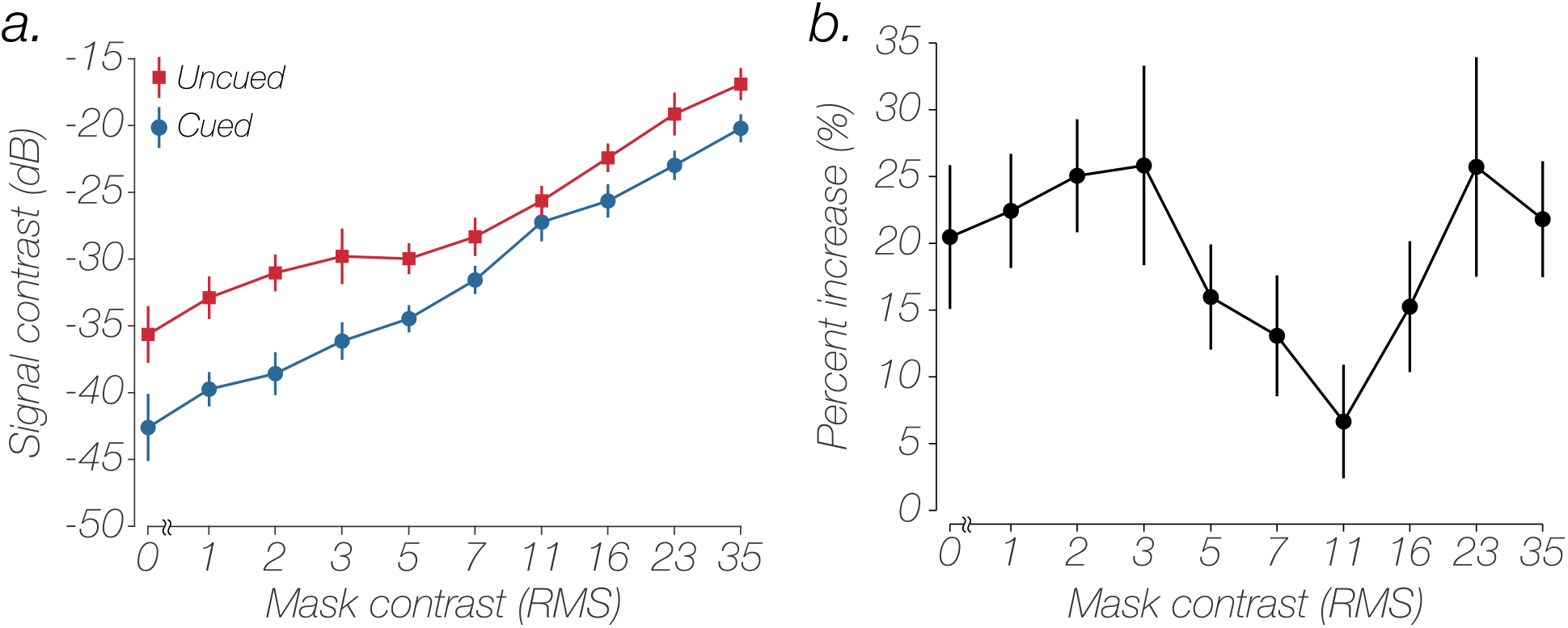
Average Perceptual Thresholds and Cuing Effect. **A.** Average perceptual thresholds across increasing levels of noise and cue presence (N=11). The red curve represents thresholds in the absence of the auditory cue, whereas the blue curve represents thresholds under the presence of the auditory cue. Thresholds in the cue condition are enhanced across all levels of noise. **B.** Average improvement in contrast sensitivity between attentional conditions, expressed as a percent increase between the cued and uncued condition. Error bars represent SEM.

**Figure 4.**
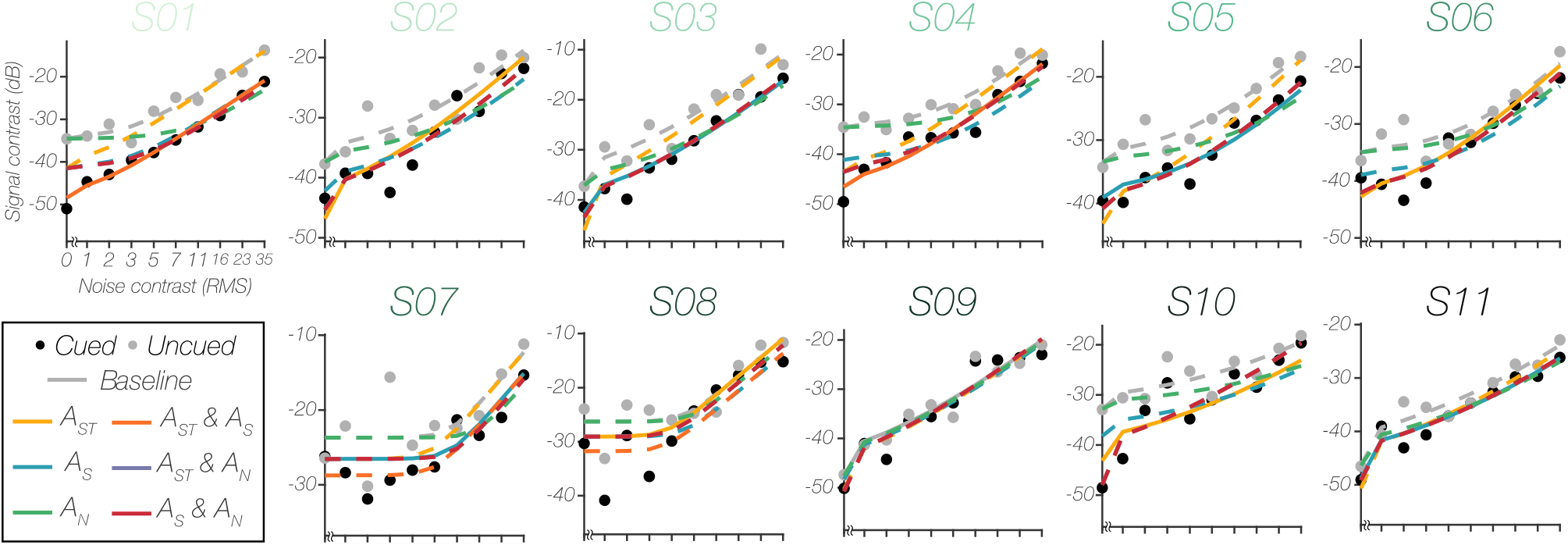
Model Fitting Results. Each plot represents an individual subject’s signal contrast threshold data for each noise contrast level (gray and black dots) and each variant of the modified normalization model of attention fit to the data from the cued condition (colored lines). Solid lines in each plot represent the winning model according to the lowest ΔAICc value for that subject. Baseline is an absence of attention coefficients/mechanisms fit to the data from the cued condition. A_ST_ represents stimulus enhancement, A_S_ represent signal enhancement, and A_N_ represents external noise exclusion. Subplot titles are color-coded to match the individual model comparison results presented in Figure 5.

**Figure 5.**
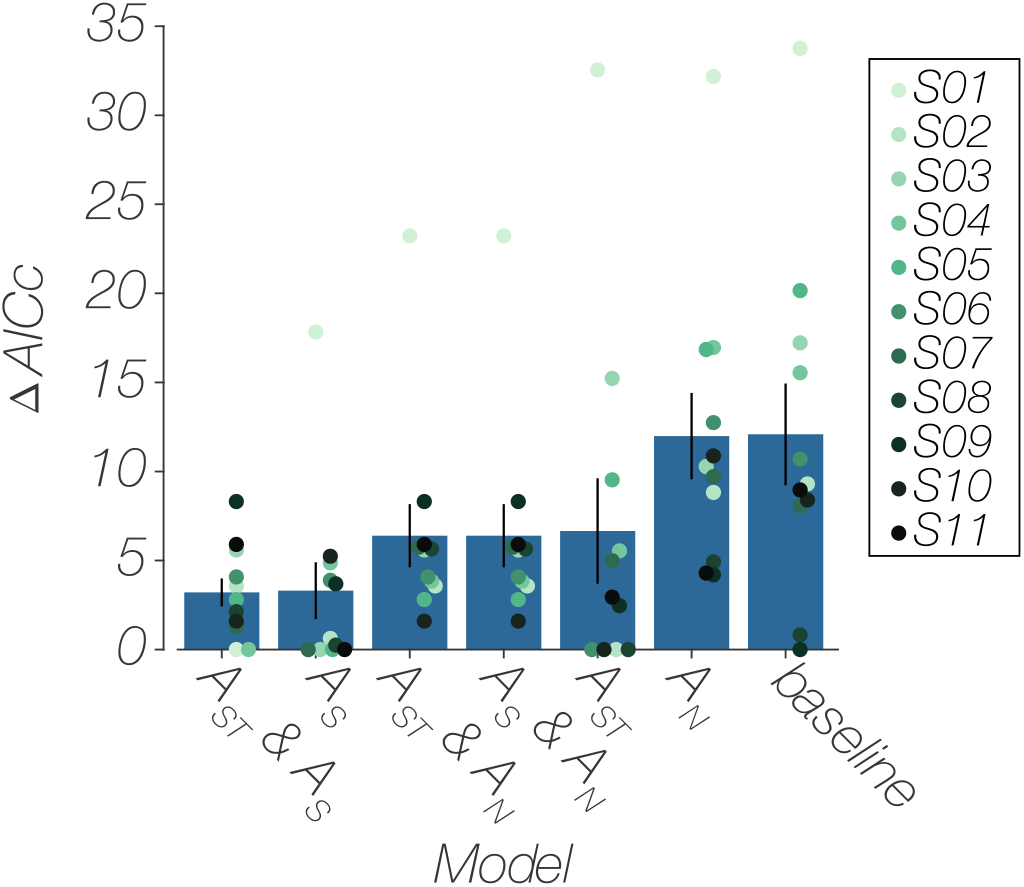
Model Comparison Results. Average fitting results for each attention mechanism (n=11). Individual subject points are jittered horizontally for better visualization. Baseline is an absence of attention coefficients/mechanisms fit to the data from the cued condition. A_ST_ represents stimulus enhancement, A_S_ represent signal enhancement, and A_N_ represents external noise exclusion. A combination of stimulus enhancement and signal enhancement had the lowest ΔAICc value on average, while signal enhancement alone closely tailed this result, suggesting some form of signal enhancement as the winning mechanism. Error bars represent SEM.

### The cueing effect cannot be explained solely by a reduction in temporal uncertainty

Our modeling results suggest that temporal attention improves fine-orientation discrimination through a combination of stimulus enhancement and signal enhancement. However, another possibility is that the temporal cue improved performance by reducing uncertainty about the moment at which the target grating appeared (Pelli 1985). Because the temporal cue in the cued condition perfectly predicted when the grating would appear, the cue may have instead improved performance by enabling participants to disregard irrelevant moments in time. Indeed, some attentional benefits have been shown to be attributed to a reduction in uncertainty, particularly with spatial attention (e.g. Gould, Wolfgang, and Smith 2007; Solomon, Lavie, and Morgan 1997). To test this uncertainty reduction account, we conducted a control experiment in which we asked participants to detect the presence or absence of a grating in each condition. We reasoned that if participants were uncertain about when the target grating (the signal) appeared, they would perform poorly in the detection task (Carrasco, Penpeci-Talgar, and Eckstein 2000). Moreover, if the temporal cue reduced temporal uncertainty, then the cue should improve detection performance.

Six participants (three from the main study and three additional participants) completed the detection task (see Methods, Participants). Figure 6a shows the signal contrast thresholds for these observers in the fine-orientation discrimination task across the 5 noise levels used in the detection task (subsampled from the 10 levels used in the main experiment, see Methods, Control experiment). As in the main study, we found that signal contrast thresholds increase with noise contrast, *F*(4,50) = 40.43, *p* < 0.001, and were lower in the cued condition than in the uncued condition, *F*(1,50) = 9.62, *p* = 0.0032. Furthermore, there was no interaction between noise level and cue condition, *F*(4,50) = 0.13, *p* = 0.9722, such that the size of the cueing effect did not scale with noise contrast, as was the case in our main experiment.

**Figure 6.**
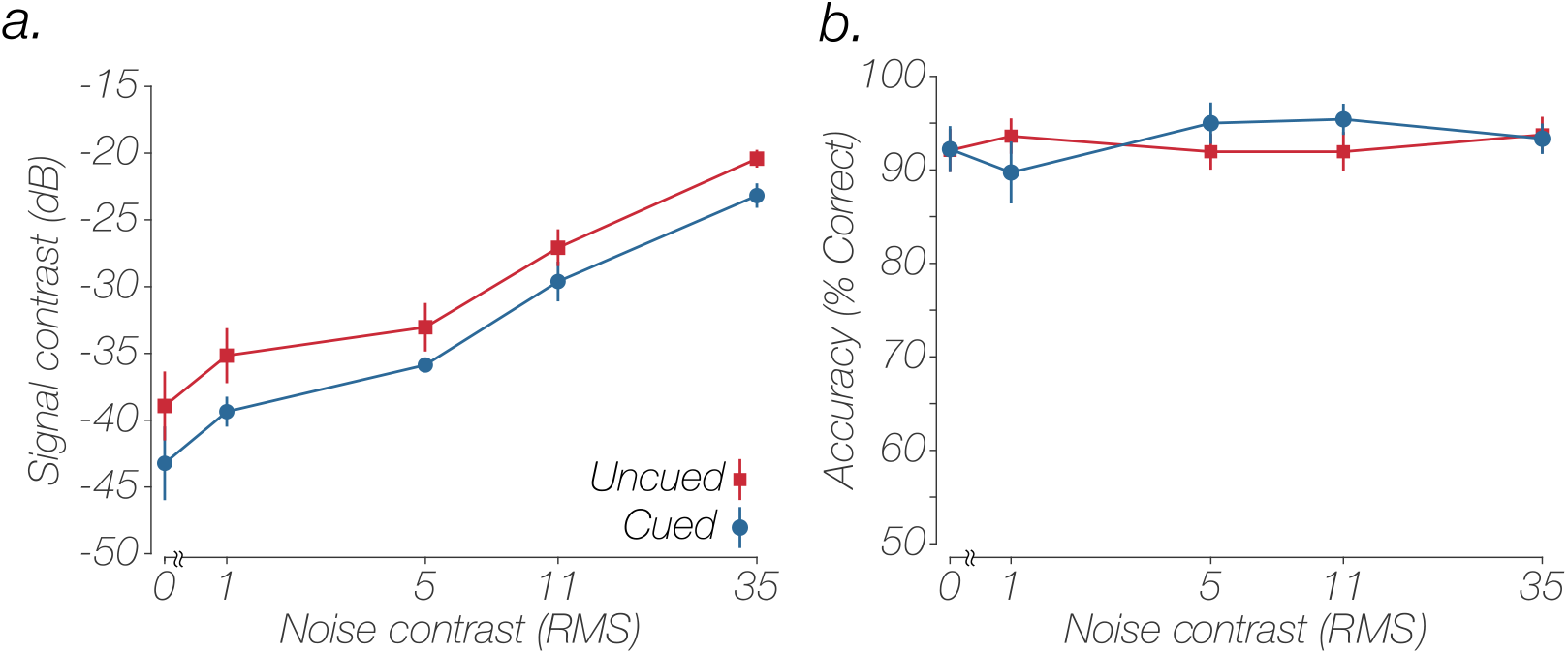
Cueing Did Not Improve Detection of Target Gratings. A. Average perceptual thresholds across increasing levels of noise and cue presence in the fine-orientation discrimination task (n=6). The red curve represents thresholds in the absence of the auditory cue, whereas the blue curve represents thresholds under the presence of the auditory cue. Thresholds in the cue condition are enhanced across all levels of noise. B. Average accuracy (n=6) in the detection task across noise mask levels and cue conditions, given signal contrasts from the fine-orientation discrimination at each noise level. There was no effect of the cue on detectability, across all noise levels, ruling out an uncertainty account for our results. Error bars represent SEM.

In the detection task, we fixed the contrast of the grating in each condition to the signal contrast thresholds estimated in the fine orientation discrimination task for each participant. Importantly, we found that detection accuracy (Figure 6b) was high in both the cued and uncued conditions (cued: 93.14% ± 2.25%; uncued: 92.67% ± 2.03%). We conducted a factorial ANOVA to test for main effects of cue (temporal uncertainty) on accuracy. Critically, we found that the temporal cue did not improve detection performance (main effect of cue: F(1,110)=0.12, *p*=0.7337; main effect of noise: F(4,110)=0.36, *p*=0.8383; interaction: F(4,110)=0.93, *p*=0.4504). These results show that the target gratings were easily detected, and the temporal cue did not improve detection performance, suggesting that temporal uncertainty did not drive the observed cuing effect in the discrimination task.

## Discussion

To date, it has remained unclear whether temporal attention increases gain for all aspects of a stimulus (stimulus enhancement) or selectively increases gain for target features (signal enhancement) to improve perception (Nobre, Correa, and Coull 2007; Nobre and Rohenkohl 2014). In this study, we parametrically varied the contrast of a noise mask — an approach used to investigate mechanisms of spatial attention (Lu and Dosher 2000) — to tease apart the mechanisms of temporal attention under a normalization framework. We found that an auditory temporal cue reduced contrast thresholds in an orientation discrimination task across all levels of external noise (Figure 3). Our modeling results revealed that this effect was best described by a combination of signal enhancement and stimulus enhancement, with signal enhancement alone a close runner up (Figure 5). Therefore, our results provide evidence against the possibility that temporal attention improves perception solely by increasing visual gain in a non-selective manner (i.e., stimulus enhancement). Instead, our results suggest that temporal attention selectively increases gain for a target feature in addition to increasing gain in general (signal enhancement and stimulus enhancement).

In a control experiment, we considered whether our cueing effect could be accounted for by a reduction of temporal uncertainty, such that observers were better able to exclude irrelevant moments in time from their decision in the cued condition than in the uncued condition. To do so, we asked observers to perform a detection task, reporting the presence or absence of a target grating (with target contrast set to the threshold measured in the discrimination task). We found that detection accuracy was high, and that our temporal cue did not improve detection performance (Figure 6). Therefore, it is unlikely that the cue reduced temporal uncertainty because the target gratings were readily detected in both the cued and uncued conditions. However, we must acknowledge that uncertainty reduction may nevertheless contribute to the cueing benefit in orientation discrimination. Even though the target gratings were suprathreshold in both conditions, it is possible that observers were still better able to exclude irrelevant moments in time for decisions in the cued condition than in the uncued condition.

One possibility as to why we observed a combination of signal enhancement and stimulus enhancement is because our temporal cue engaged multiple processes. We manipulated temporal attention using a temporal orienting auditory cue that swept in pitch over 500-ms, preceding the target stimulus. Temporal orienting cues are commonly used in studies of temporal attention (Griffin, Miniussi, and Nobre 2001; Coull et al. 2000; Nobre 2001; Correa, Lupiáñez, et al. 2006; Correa et al. 2004). However, while temporal orienting cues allow observers to voluntarily deploy endogenous temporal attention, the cue itself may also trigger a reflexive increase in alertness or arousal (Weinbach and Henik 2012). Recent work has begun to tease apart the effects of endogenous and exogenous (i.e., reflexive) temporal attention (Lawrence and Klein 2013), with some studies suggesting that endogenous and exogenous temporal attention have dissociable effects on perception (McCormick et al. 2018; Rohenkohl, Coull, and Nobre 2011). One possibility in our study is that exogenous temporal attention was responsible for the overall increase in gain and endogenous temporal attention was responsible for selectively increasing gain for target features. However, further work is needed to test whether signal enhancement and stimulus enhancement effects are specifically linked with endogenous or exogenous orienting of temporal attention, under a masking paradigm and normalization framework.

In conclusion, we used a masking paradigm and a normalization framework to tease apart the mechanism subserving temporal attention. Under this framework, temporal attention can improve visual sensitivity through stimulus enhancement — amplifying everything attention is directed towards, signal enhancement — selectively enhancing *just* the signal and leaving irrelevant noise untouched, or external noise exclusion — leaving the signal untouched and actively suppressing irrelevant noise. Because previous studies of temporal attention have not manipulated external noise, it has remained unclear whether temporal attention increases gain for all aspects of a stimulus (stimulus enhancement) or selectively increases gain for target features (signal enhancement) to improve perception. Here, we found that temporal attention recruits both signal enhancement and stimulus enhancement, such that temporal attention selectively enhances the processing of target features as well as producing a general increase in gain.

## Footnote

We note that the terms “stimulus enhancement” and signal enhancement” have been used interchangeably in past work to refer to what we call stimulus enhancement, a wholesale increase in gain that will amplify target features and noise (e.g. Lu and Dosher 1998; Dosher and Lu 2000; Ling and Carrasco 2006). Dosher and Lu (2000) rightly noted that stimulus enhancement might be the better term when both the signal and noise are being modulated. In this paper, we follow their lead. Thus, we use stimulus enhancement to refer to a wholesale increase in gain, and signal enhancement to refer to an increase in gain for target features.

## Acknowledgements

We thank Dr. Rachel Denison, her lab, and the Ling Lab for their invaluable feedback. This research was funded by National Institutes of Health Grant EY028163 to S. Ling.

